# Characterization of the ligand binding pocket of the virulence regulator Rns, a Member of the AraC/XylS family of transcription factors

**DOI:** 10.1101/2025.01.14.633009

**Authors:** Jessica D Tolbert, Kacey M Talbot, F. Jon Kull, George P Munson, Charles R Midgett

**Affiliations:** Department of Chemistry, Dartmouth College, Hanover, NH 03755; Department of Microbiology & Immunology, University of Miami, Leonard M. Miller School of Medicine, Miami, FL 33101

**Author notes:** Corresponding authors: George P Munson;, Charles R Midgett. University of North Carolina at Chapel Hill, Chapel Hill, NC 27514. These authors contributed equally to this work (Jessica D Tolbert, Kacey M Talbot).

## Abstract

Diarrheal disease caused by Gram-negative enteric pathogens such as enterotoxigenic *Escherichia coli* (ETEC), *Vibrio cholerae*, *Shigella spp.*, and, *Salmonella spp.* are a leading cause of morbidity and mortality of children; especially, in low resource nations. While progress has been made in reducing this burden, there remains a need to develop efficacious therapies.

Recently, we determined the structure of Rns, a member of the AraC/XylS family that regulates the expression of pili and other virulence factors in ETEC. The structure revealed decanoic acid bound between the N- and C-terminal domains. To test the hypothesis that bound decanoic acid directly inhibits Rns, we used its structure to identify residues that are necessary for ligand binding. Removal of the positive side chains of R75 and H20 rendered Rns insensitive to fatty acid inhibition. Additionally, mutations designed to occlude decanoic acid binding also produced a variant that was fatty acid insensitive. We also observed that this variant is structurally more flexible than wildtype Rns with decanoic acid; suggesting that bound fatty acid contributes to structural rigidity.

These studies precisely demonstrate Rns binding pocket residues critical for binding fatty acids and inhibition of DNA binding. This supports our hypothesis that fatty acids must bind in the binding pocket to inhibit AraC regulators. Further work by us and others suggests inhibition of AraC virulence regulators by fatty acids is a common paradigm among many bacterial pathogens. Therefore, understanding the molecular basis of this inhibition lays the groundwork for the development of small molecule therapeutics targeting enteric disease.

## Introduction

Diarrheal disease is the second leading cause of death in children worldwide, leading to an estimated half a million deaths annually(WHO 2024). Enterotoxigenic *E. coli* (ETEC) accounts for close to fifty thousand deaths of children under five (WHO 2024). Survivors of repeated ETEC infections may suffer malnutrition during their critical developmental years and suffer comorbidities such as stunting and long-term cognitive impairment(Anderson et al. 2019; Fleckenstein and Kuhlmann 2019). Much effort has been expended to develop a vaccine against ETEC. However, vaccine development has mainly focused on pilins and those efforts have been stymied by the antigenic diversity of pilins and suboptimal long-term immunity (Fleckenstein 2021). This, combined with a rise in multidrug resistant *E. coli* strains, highlights the need for alternative therapeutic solutions (Cruz-Córdova et al. 2014; Eltai et al. 2020; Joffré and Rojas 2020). One such approach is to focus on pathways specific to the pathogenic process which should decrease the selection for antibiotic resistance (Ellermann and Sperandio 2020).

ETEC and other enteric pathogens possess transcriptional regulators belonging to the AraC/XylS family that directly regulate the expression of genes involved in pathogenesis, including attachment factors, the diarrheal causing toxins, and other virulence factors such as CexE, AcfA, and AcfD, (Caron et al. 1989; DiRita et al. 1991; Withey and DiRita 2005; Bodero et al. 2007; Rivas et al. 2020).Like other canonical AraC/XylS family members, these virulence regulators are defined by a structurally conserved DNA binding domain consisting of two helix-turn-helix motifs (Gallegos et al. 1997) and an amino-terminal domain that participates in dimerization and/or ligand interactions (Soisson et al. 1997). Rns, a canonical AraC/XylS family member from ETEC, positively regulates its own expression and activates the expression of adhesive pili (Caron et al. 1989; Caron and Scott 1990; Haan et al. 1991; Munson and Scott 2000; Bodero et al. 2008; Bodero and Munson 2016). Without the expression of these pili adherence to the host epithelium is impaired and diarrheal disease is attenuated (Evans et al. 1978). Moreover, Rns is an attractive therapeutic target because it is well conserved across ETEC strains even though the pilins that it regulates are not. Rns also activates the expression of CexE, an outer membrane lipoprotein and virulence factor, as well as its secretion system (Pilonieta et al. 2007; Belmont-Monroy et al. 2020; Rivas et al. 2020; Icke et al. 2021). It also represses the expression of *nlpA*, a gene that has been implicated in the biogenesis of outer membrane vesicle (McBroom et al. 2006; Bodero et al. 2007). Thus, a clear understanding of the interactions between Rns and inhibitory could lead to new strategies to combat ETEC infections.

Until recently there was no evidence that Rns was directly modulated by effectors, despite the known exogenous ligand regulation of many other AraC/XylS members (Basturea et al. 2008; Kolin et al. 2008; Schleif 2010). Previous work with other AraC members like ToxT in *V. cholerae* and work from other groups on regulators from Salmonella (HilD, HilC, RtsA) and *Shigella flexneri* VirF revealed the inhibitory effects of a range of long chain fatty acids on regulator activity (Lowden et al. 2009; Childers et al. 2011; Golubeva et al. 2016; Bosire et al. 2020; Chowdhury et al. 2021; Trirocco et al. 2023). This, combined with our work on Rns, has challenged the paradigm that the ability of AraC/XylS virulence regulators to bind DNA is regulated only by protein-protein interactions, showing DNA binding is also regulated by small molecules (Cortés-Avalos et al. 2021).

The first crystal structure of Rns revealed a binding pocket between the N- and C-terminal domains occupied by the 10-carbon saturated fatty acid, decanoic acid (Midgett et al. 2021) (**Fig 1**). To investigate the biological significance of our crystallographic discovery, we cultured ETEC strains with decanoic acid concentrations from 20 µM to 1.2 mM (Midgett et al. 2021). Those experiments revealed that the expression of pilins and CexE decreased with increasing concentrations of decanoic acid and that expression of pilins was undetectable in the presence of ∼300 µM decanoic acid. We further showed that decanoic acid inhibited the expression of β-galactosidase in a Lac reporter strain in which the Rns-dependent CS3 pilin promoter drives the expression of *lacZ*. These effects were specific for proteins within the Rns regulon because exogenous decanoic acid had negligible effect on the expression of Rns-independent proteins such as flagellin. Although these results did not reveal a molecular mechanism, they suggest that the activity of Rns is abolished when decanoic acid occupies its binding pocket.

**Figure 1.**
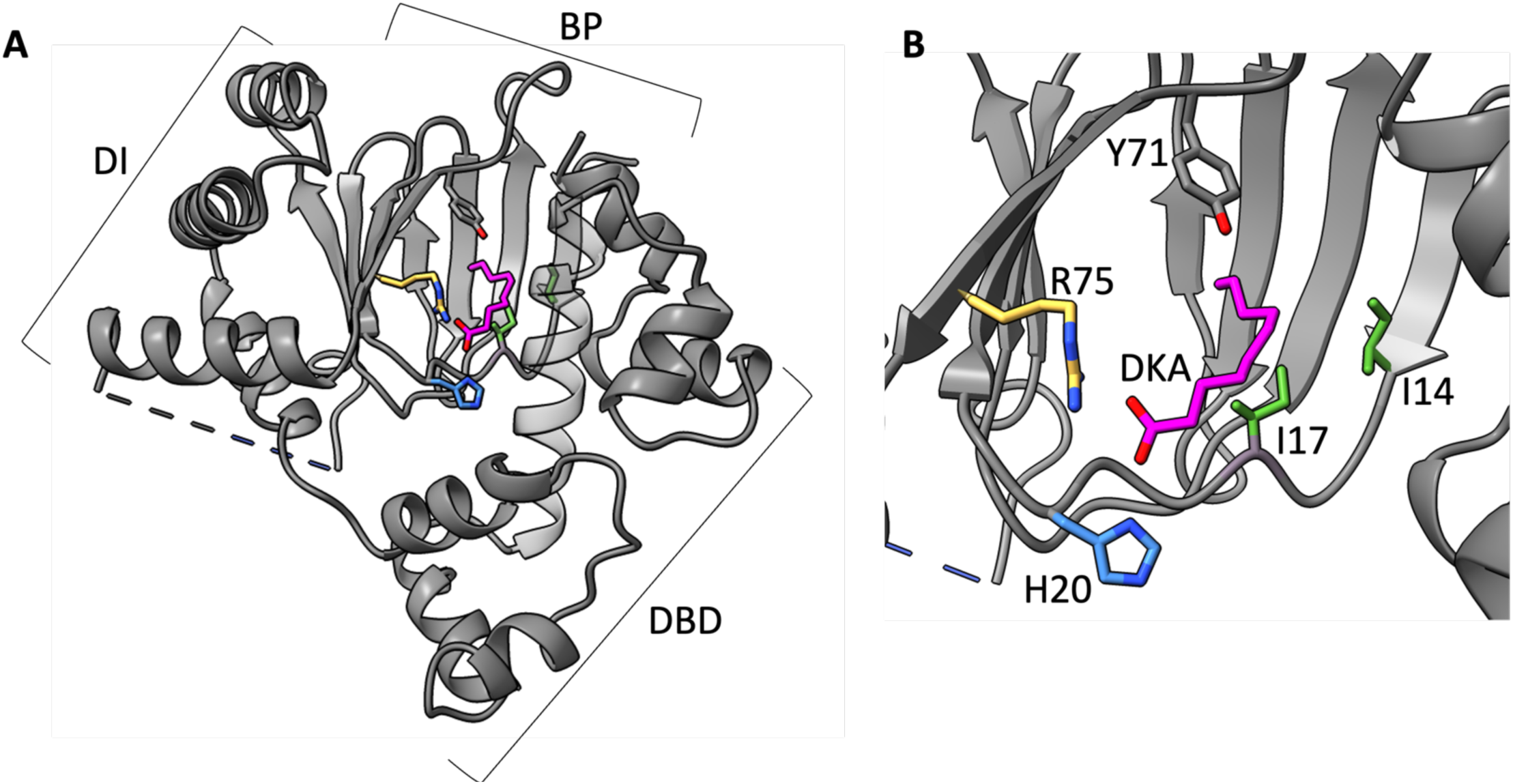
The structure of Rns with bound fatty acid. **(A)** Rns (PDB ID: **6XIU)** bound to saturated, decanoic acid. Depicted are the three domains: dimerization interface (DI), binding pocket (BP), and DNA binding domain (DBD). (**B)** Close up of the pocket showing R75 in yellow, H20 in blue, I14 and I17 in green, as well as Y71 in gray. Residue Y71 is used as a reference for all structures as an unvarying pocket element.

In this study we identify the amino acid side chains in the Rns binding pocket are critical for both fatty acid binding and inhibition of Rns. Specifically, we focused on the role of residues that stabilize the negative charge of the fatty acid carboxylate head group, as well as those positioned to block fatty acid binding following the introduction of bulkier side chains. Altering the electrostatic and volume environment of the pocket decreased the response of Rns to fatty acid, supporting our hypothesis that direct binding of fatty acid to the Rns pocket is critical for its inhibitory response. These results provide evidence that AraC family virulence regulators share a common inhibitory mechanism, laying the groundwork for future development of antivirulence therapeutics against a wide array of enteric pathogens.

## Results

### Decanoic acid interferes with Rns DNA binding

As discussed above, we have previously established that Rns binds decanoic acid and that decanoic acid inhibits its activity as a transcriptional regulator (Midgett et al. 2021). To elucidate the molecular mechanism(s) of the latter observation we considered the possibility that decanoic acid binding increases the turnover/degradation of Rns *in situ*. Since there are no validated antibodies for the detection of native Rns, we evaluated decanoic acid-dependent turnover of epitope tagged Rns in K-12 Lac reporter strains (**Fig 2A**). Although we have previously shown that 1-3 mM decanoic acid is sufficient to inhibit Rns activity in these strains, we did not observe a dramatic decrease in the level of Rns even at 5 mM decanoic acid. Likewise, a time course in which *de novo* translation was inhibited in an ETEC strain found little difference between decanoic acid treated and DMSO control cultures (**Fig. 2B**). Although qualitative, these results suggest decanoic acid binding to Rns does not appreciably affect its stability *in situ*.

**Figure 2.**
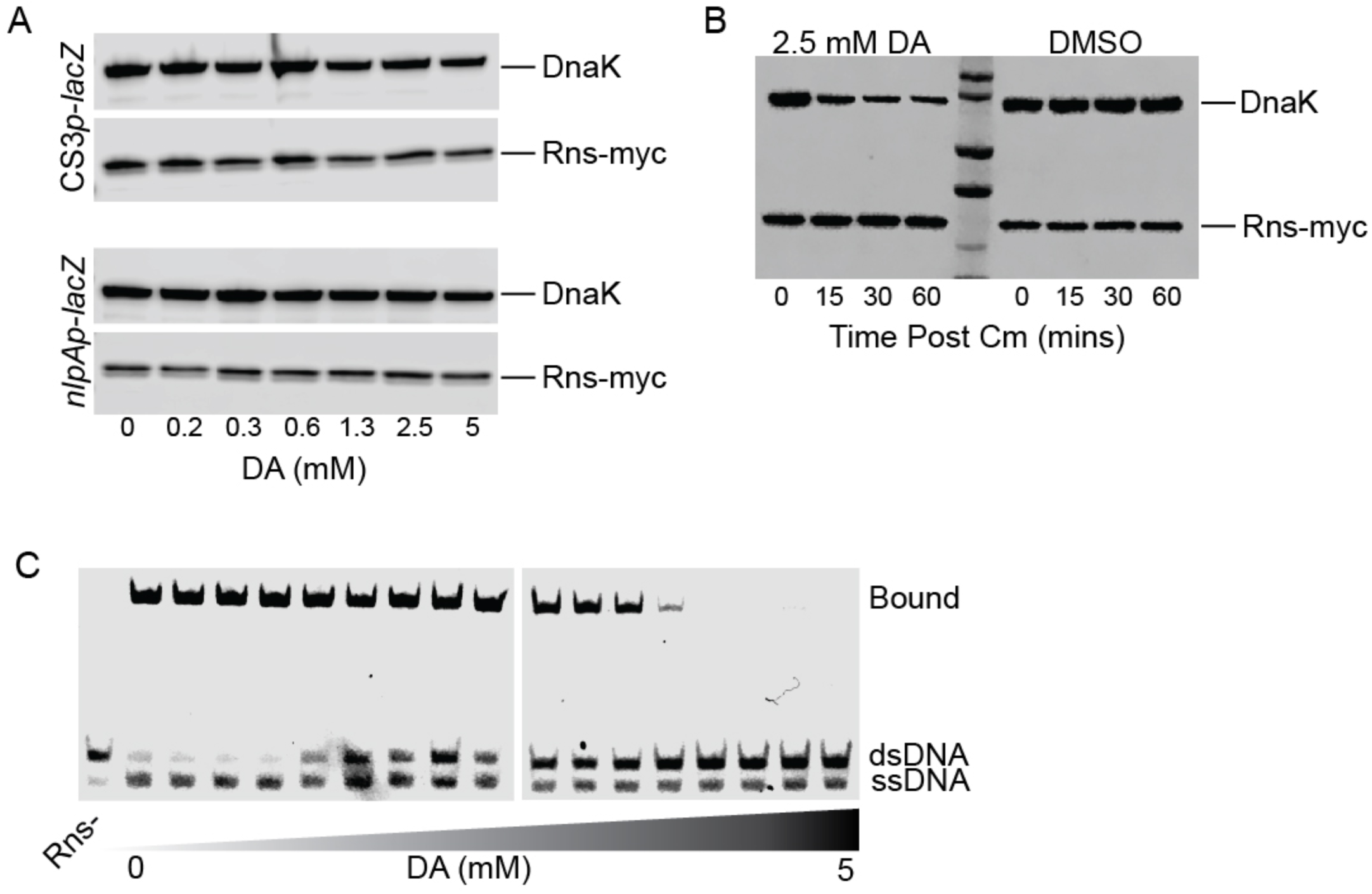
Decanoic acid does not destabilize Rns but does abolish its ability to bind DNA. (**A**) Western blots of whole cell lysates of K-12 Lac reporter strains GPM1072 and GPM1080 transformed with pGPMRns-myc. Rns-myc expression is persistent in situ relative to the DnaK loading control even at concentrations of decanoic acid that abolish its inhibitory activity. (**B**) Western blot of plasmid expressed Rns-myc in ETEC strain H10407 *rns::ka*n (GPM1236) in which translation was inhibited with 30 µg/ml chloramphenicol (Cm). The apparent turnover rate of Rns was observed to be comparable in the presence or absence of decanoic acid. (**C**) EMSA of MBP-Rns with DNA containing a prototypical Rns binding site described in Materials and Methods. Decanoic acid abolishes the ability of Rns to bind the DNA duplex in a dose-dependent manner.

As a transcription factor, the activity of Rns is dependent upon its ability to bind DNA. Thus, in the absence of apparent decanoic acid-dependent turnover of Rns we considered the possibility that fatty acid binding to the pocket abolishes DNA binding. Consistent with this hypothesis, we observed that decanoic acid did abolish the ability of Rns to bind a DNA duplex carrying a prototypical Rns binding site (**Fig. 2C**), indicating decanoic acid inhibits Rns DNA binding by interacting with the protein.

### Electrostatic interactions contribute to occupancy of the decanoic acid binding pocket

Our previous work with Rns-decanoic acid cocrystals showed the positively charged side chains of H20 and R75 are positioned ∼3.0 Å from the carboxylate head group of decanoic acid, a distance consistent with electrostatic stabilization of ligand binding **(Fig 1)**. To evaluate if decanoic acid binding is dependent upon the electrostatic interactions, we determined the crystal structures of Rns–H20A, – R75A, and –H20A/R75A to 2.9 Å, 2.35 Å, and 2.4 Å resolution respectively. Overall, the structures of –H20A, –R75A, and –H20A/R75A were very similar to wildtype. However, in each of the three variants the binding pocket showed no electron density that would be consistent with a bound fatty acid (**Fig. 3**). These results suggest that electrostatic interactions between the carboxylate head group of decanoic acid and the side chains of H20 and R75 are important for fatty acid binding.

**Fig 3.**
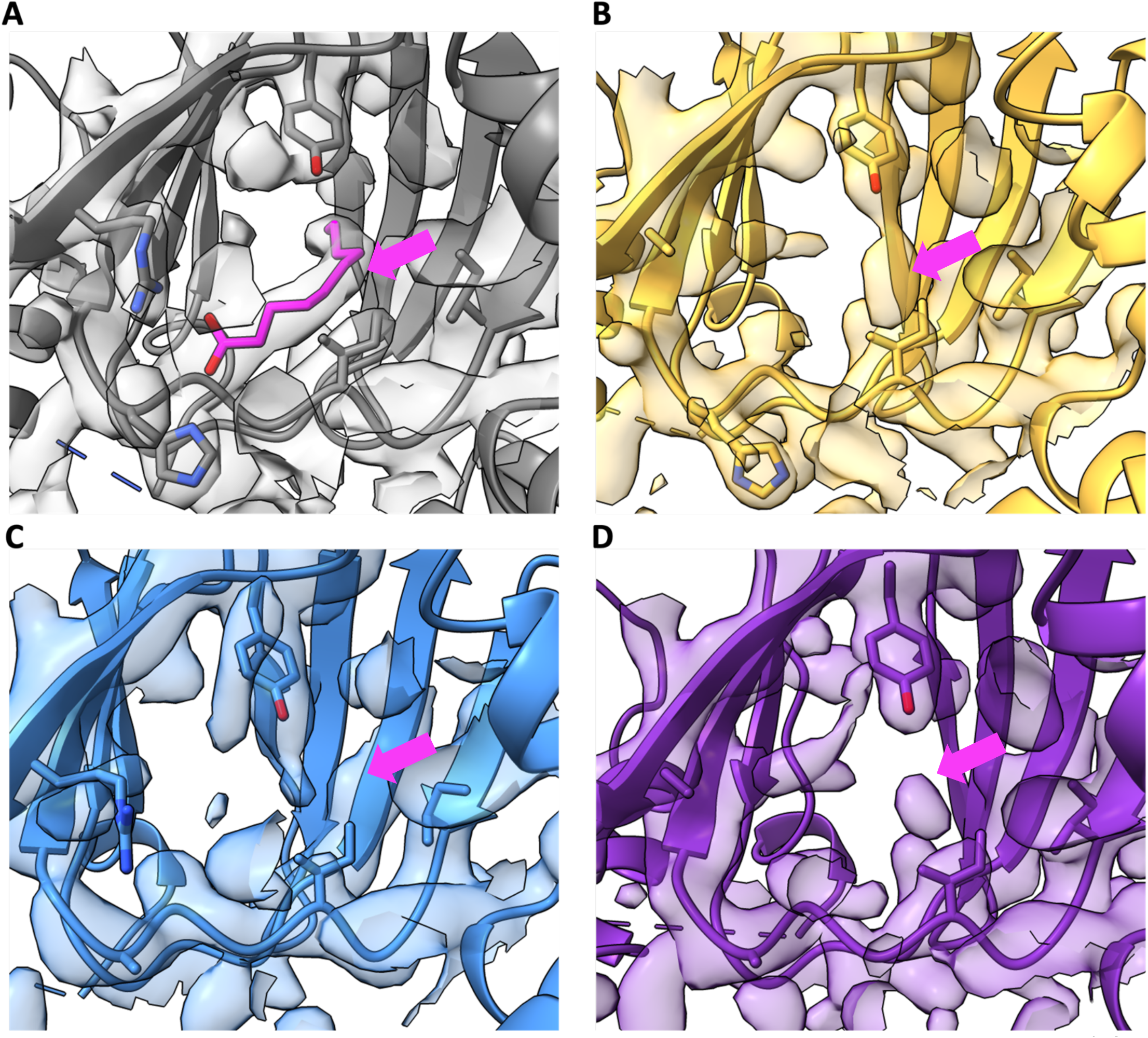
Rns pocket mutants abolish previously observed electrostatic interaction potential. (**A)** WT Rns (grey) with bound decanoic acid. Variants (**B**) R75A (yellow, PDB ID: 8VRQ), (**C**) H20A (blue, PDB ID: 9CA5) and (**D**) H20A/R75A (purple, PDB ID: 8VST) do not contain electron density consistent with bound ligand. Y71 is visible in all the structures.

To further validate our crystallographic findings, we evaluated the activity and decanoic acid responsiveness of WT Rns and the three variants. In the absence of decanoic acid, the single and double alanine variants were active as determined by Rns-dependent expression of β-galactosidase in a CS3p–Lac reporter strain. Relative to the DMSO control, the activity of WT Rns decreased by 90% with 5 mM decanoic acid (**Fig. 4**). In contrast, the activity of Rns–R75A and –H20A/R75A was inhibited by 16% and 8% respectively. Rns–H20A displayed an intermediate effect at 68% inhibition by decanoic acid.

**Figure 4.**
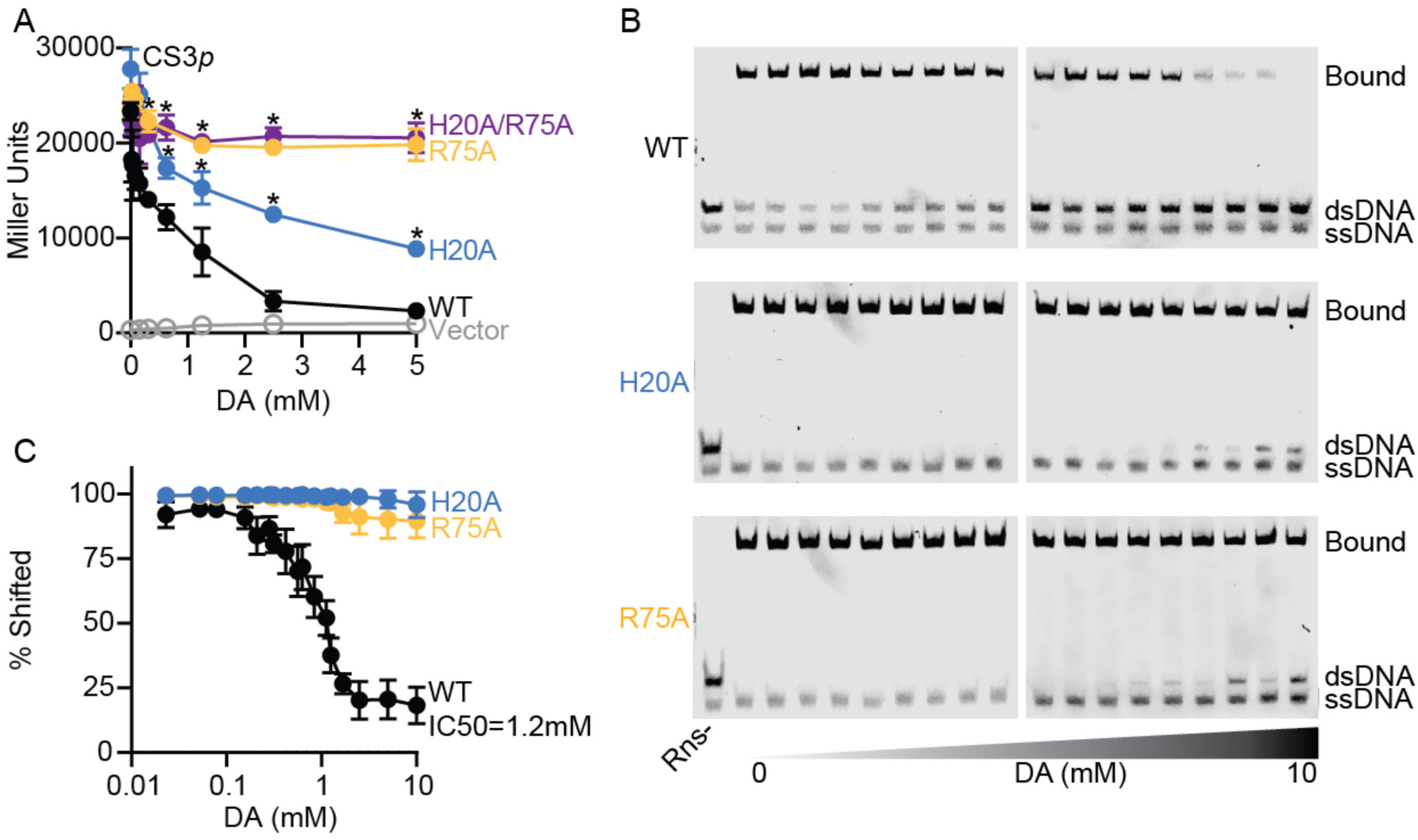
Decanoic acid inhibition is dependent on interactions between decanoic acid and Rns residues H20 and R75. (**A**) β-galactosidase reporter assays of Rns-WT, H20A, R75A, and H20A/R75A in the presence of decanoic acid in 0.4% (v/v) DMSO. Data is shown as the mean response ± SD, \**P* < 0.05 by Student’s t-test compared to WT activity. *n* = 3. (**B**) Representative EMSAs showing the effect of decanoic acid on 25 nM MBP-Rns binding to 5 nM DNA with a prototypical Rns binding site. (**C**) Quantification of *n* = 3 EMSAs using Licor Image Studio Lite. Data is shown as percent shifted (shifted band density divided by density of entire lane) ± SD. Decanoic acid inhibits DNA binding of Rns-WT with an IC50 of 1.2 mM and has no effect on DNA binding for Rns-H20A and R75A.

In EMSAs, the IC50 for decanoic acid with WT Rns was determined to be 1.15 mM. The IC50 is estimated to exceed 10 mM decanoic acid for Rns–H20A and –R75A but could not be quantified due to the fatty acid’s inability to fully dislodge the variants from DNA (**Fig. 4**). The critical micelle concentration of decanoic acid also precluded titrations beyond ca. 15 mM. Interestingly, Rns–H20A appears to be less sensitive to decanoic acid by EMSA than Lac reporter assays. We attribute this inconsistency to intrinsic differences between the *in vitro* and *in situ* assays. These results indicate that the side chains of both H20 and R75 interact with the carboxylate head of decanoic acid and that fatty acid binding to the pocket abolishes the ability of Rns to bind DNA.

### Occlusion of the fatty acid binding pocket

To further investigate of the structural requirements for ligand binding and response, we introduced mutations to sterically occlude ligand binding. Specifically, residues I14 and I17, which line the hydrophobic region of the pocket, were mutagenized to phenylalanine or tyrosine. As expected from our analysis, Rns–I14F/I17F was far less sensitive to decanoic acid inhibition than WT Rns; 25% vs. 75% max inhibitory activity (**Fig. 5**). As with the H20A and R75A variants, the I14F/I17F is also estimated to have an IC50 of >10 mM decanoic acid by EMSA. Although the I14Y/I17Y also appeared to be largely insensitive to decanoic acid in the Lac reporter strain, it was not investigated further due to low overall activity relative to WT Rns and the I14F/I17F variant. To confirm that decanoic acid insensitivity was due to lack of fatty acid binding, we crystallized the I14F/I17F variant and solved its structure to 3.0 Å. As with the mutations that abolish electrostatic interactions with the carboxylate head group of decanoic acid, we observed no fatty acid like electron density in the altered binding pocket (**Fig 6A**). This is likely because the bulky side chain phenyl groups of F14 and F17 occludes the position occupied by the aliphatic chain of decanoic acid in the WT structure (**Fig. 6B**). Unlike the electrostatic mutants, the structure of I14F/I17F differed from that of WT Rns. The side chains of both H20 and R75 moved into the pocket, with the imidazole ring of H20 occupying part of the space where the carboxylate head group of the decanoic acid ligand is in the WT structure (**Fig. 6C**).

**Figure 5.**
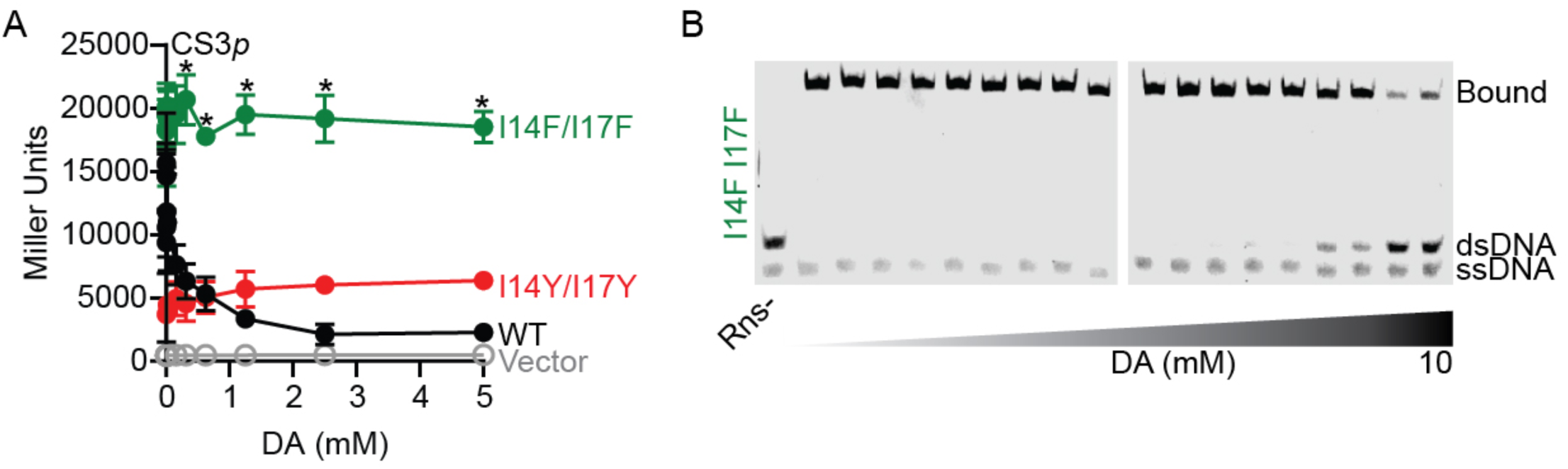
Rns-I14F/I17F is not inhibited by decanoic acid. (**A**) β-galactosidase assays of Rns-WT, I14F/I17F, and I14Y/I17Y in the CS3p Lac reporter strain. Decanoic acid inhibits Rns-WT but does not have an effect on Rns-I14F/I17F or I14Y/I17Y. Data is given as the mean response ± SD, * *P* < 0.05 by Student’s *t*-test compared to WT activity. *n* = 3. (b). (**B**) EMSA showing the effect of decanoic acid on 25 nM MBP-Rns-I14F/I17F binding to 5 nM DNA with a prototypical Rns binding site.

**Figure 6.**
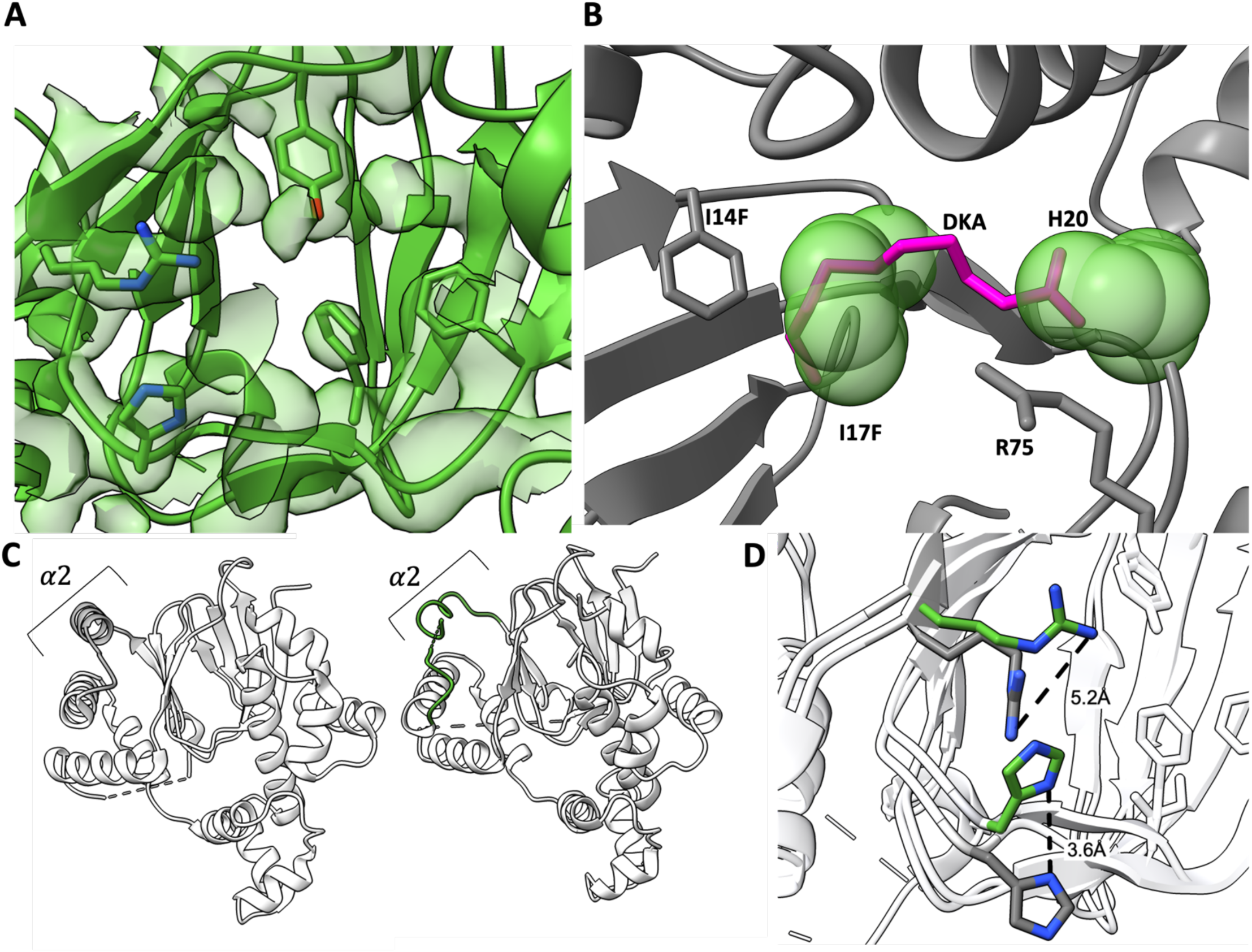
Analysis of the I14F/I17F pocket variant. (**A**) I14F/I17F (green, PDB ID: 9CA6) structure overlayed with corresponding electron density map shows no sign of bound decanoic acid. (**B**) Superposition of WT Rns and bound decanoic acid (fuchsia, showing represented van der waals radii) with the I14F/I17F structure, showing overlap of the variant side chains and the space occupied by decanoic acid. (**C**) Helix a2 (grey in WT Rns) becomes disordered in the I14F/I17F (green) variant. This change did not affect subsequent crystallization packing. (**D**) The side chain of R75 moves by 5.2 Å into the pocket of the I14F/I17F mutant when compared to WT, and the side chain of H20 moves nearly 4 Å up and toward R75 in the I14F/I17F mutant.

### Overall structural analysis

To determine if there were any structural differences between the previously published wildtype structure (Midgett et al. 2021) and the altered proteins we aligned the backbone atoms of the structures as well as compared their scaled B-factors. Overall, the structures and scaled B-factors are similar between all the structures. Aligning the back bones between the wildtype and altered protein structures gave RMSDs of 0.574 to 1.17 Å^2^. Even though the structures are similar there are two differences between the wildtype and altered proteins that we observed. First, the dimerization interface helix a2 in I14F/I17F is a coil with part of the connection to helix a3 unresolved. **(Fig. 6D)**. The second difference is all the mutant proteins show a more open pocket as measured from the long DBD helix *α*8 to β7 (residue 73, and 219). In the absence of ligand, the distance between the alpha-carbons of A73 and L219 in all mutants increasing by ∼1 Å (**Fig. 7**). Taken together, these results indicate that the altered proteins, which appear to be ligand free, are more flexible than the wildtype protein.

**Figure 7.**
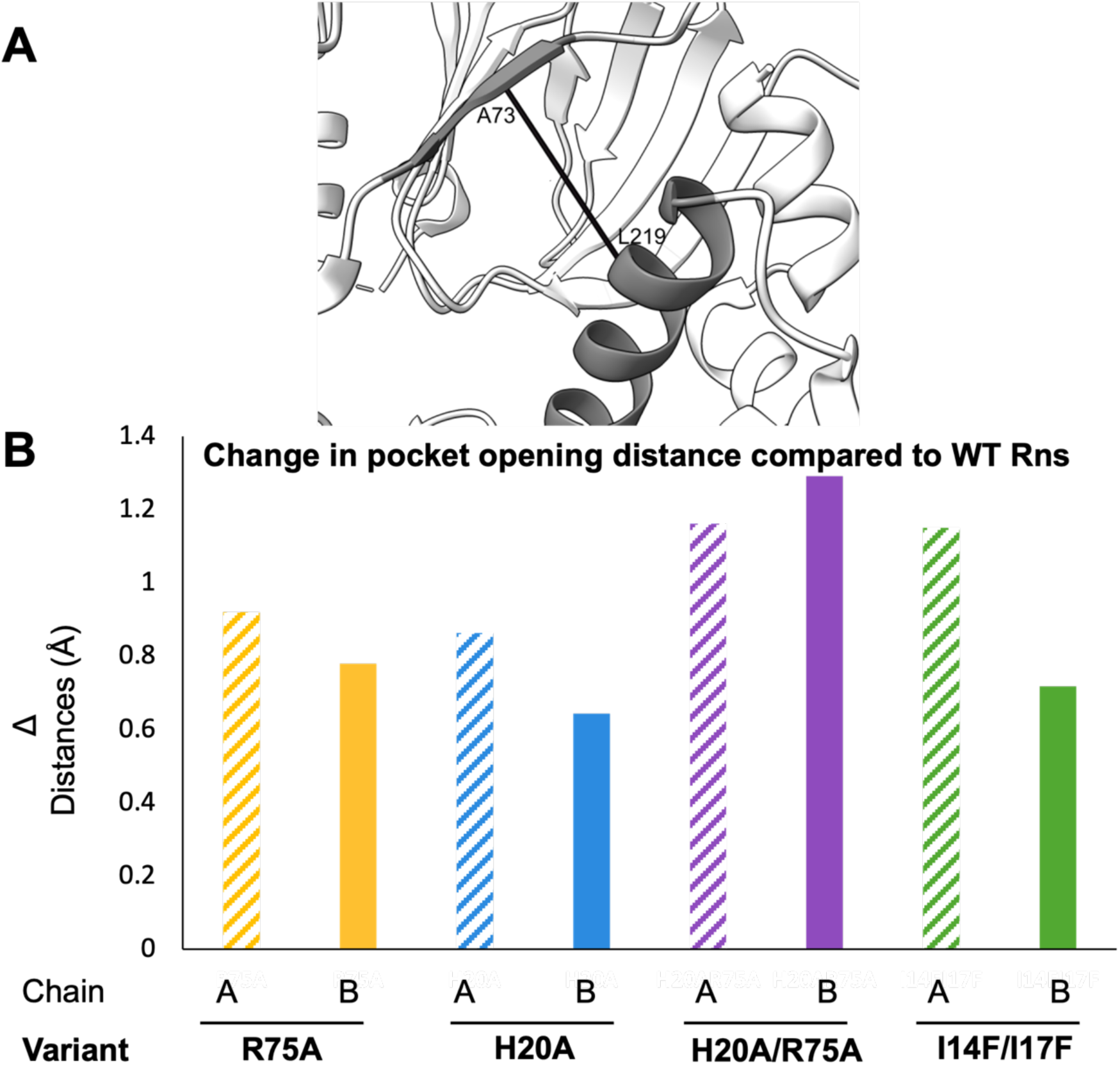
Global changes in the pocket of WT and variant Rns structures. (**A) Widest point of pocket opening.** All measurements were taken at the CA of residues A73 and L219. (**B)** Change in width compared to WT.

To determine if there were differences in protein chain flexibility between the wildtype and altered proteins the B-factors were scaled using inter-quartile scaling. Graphing the scaled Cα B-factors for the wildtype and altered proteins showed they followed a similar pattern across the structures (**Fig, S1**), with one notable exception. The Rns-I14F/I17F structure has more disorder in the first DNA binding helix compared to the other structures. This suggests changes in flexibility are reflected in the structures as seen above and not necessarily reflected in the B-factors.

### Specificity/promiscuity of the fatty acid binding pocket

Although Rns was initially crystallized with decanoic acid, we considered the possibility that other fatty acids may also occupy the pocket and inhibit the activity of Rns. Indeed, hexanoic acid (C6), octanoic (C8), and dodecanoic acid (C12) saturated fatty acids significantly inhibited the activity of Rns in our β-galactosidase assay (**Fig. 8A**). We also observed inhibition by oleic acid (C18:1), an unsaturated fatty acid. The Rns-H20A/R75A variant was found to be insensitive to the same fatty acids (**Fig. 8B**). These results suggest that the carboxylate head groups of each are coordinated by electrostatic interactions with the side chains of H20 and R75, as we observed for decanoic acid. With the exception of hexanoic acid, occlusion of the binding pocket in Rns-I14F/I17F also abolished inhibition by the fatty acids (**Fig. 8C**). Although the I14F/I17F mutation may not fully prevent binding of the aforementioned fatty acids, this mutation did decrease the effectiveness of inhibition relative to WT Rns.

**Figure 8.**
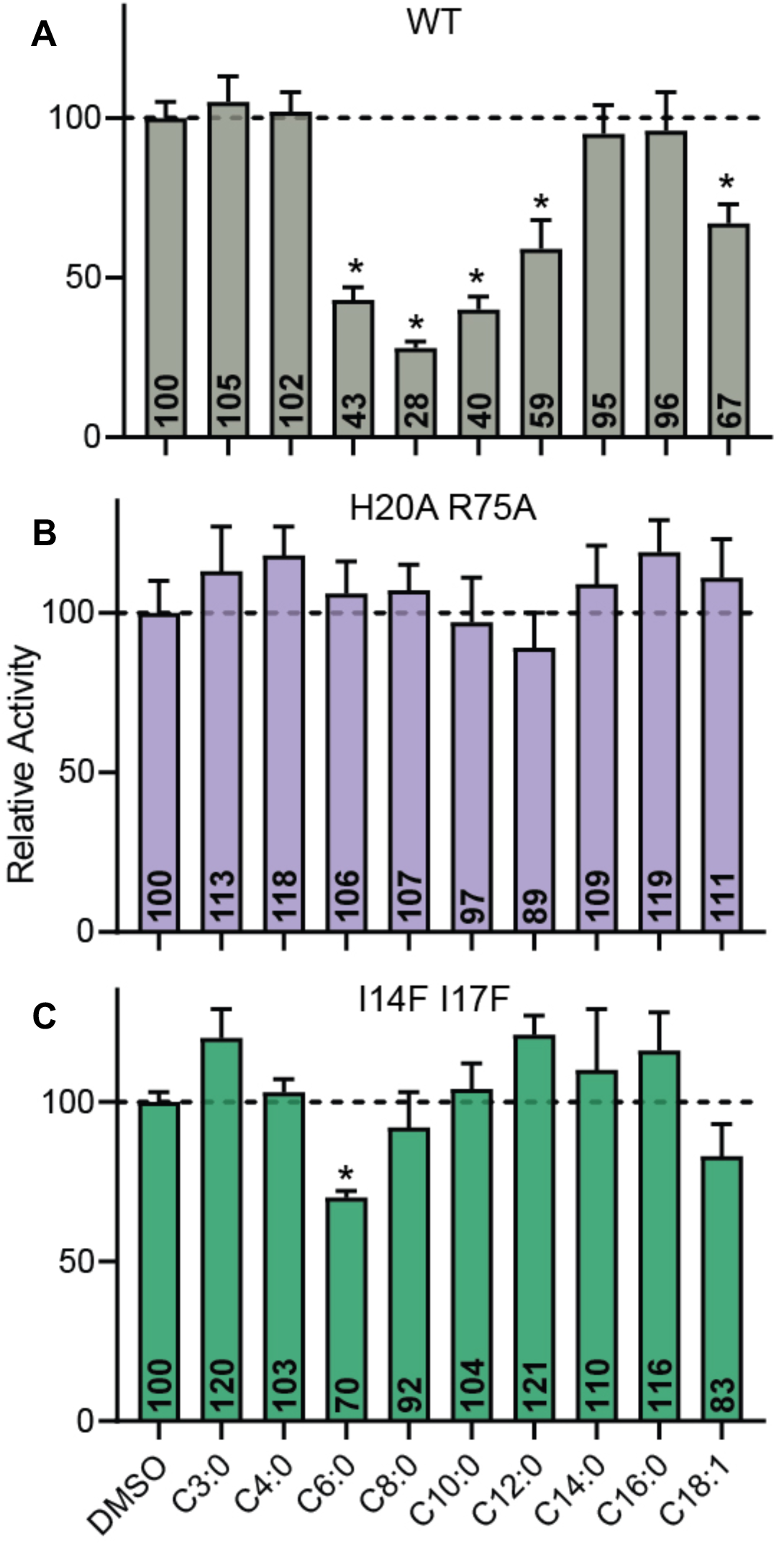
The specificity of the Rns fatty acid binding pocket. β-galactosidase assays of Rns-WT (**A**), H20A/R75A (**B**), and I14F/I17F (**C**) expressed in CS3p reporters in the presence of 3 mM (0.4% final DMSO v/v) of saturated fatty acids C3-C16 or monounsaturated fatty acid C18:1. Data is given as mean relative activity (activity normalized to DMSO x 100) ± SD, *n* = 3. * *P* < 0.002 to DMSO activity via Student’s *t*-test.

## Discussion

There is a growing body of evidence that many AraC-virulence regulators (AraC-VR) from enteric bacteria are inhibited by fatty acid ligands. Since it was found that monounsaturated long chain fatty binding to *V. cholerae* ToxT inhibits virulence gene transcription by disrupting DNA binding (Lowden et al. 2009; Childers et al. 2011), others have shown that fatty acids similarly inhibit virulence gene expression in *Salmonella* (HilD, HilC, and RtsA) (Golubeva et al. 2016; Chowdhury et al. 2021). We recently showed that fatty acid binding to Rns from ETEC also inhibits virulence gene expression and identified a number of additional AraC-family members that we hypothesized based on predicted structural similarities could be regulated in a similar manner, including FapR (*E*. *coli* ETEC), PerA (*E. Coli* EPEC), HilC (*Salmonella typhi*), VirF (*Shigella flexneri*), and VirF (*Yersinia enterocolitica*). Supporting this hypothesis, it was recently shown that VirF from *S. flexneri* is indeed regulated by medium-chain saturated and long-chain unsaturated fatty acids (Trirocco et al. 2023). Despite this evidence, data that directly shows fatty acid ligand binding in the pocket regulates activity had not been shown. In this study, we used the structure of Rns bound to decanoic acid to investigate the roles of the binding pocket residues in binding ligand, and regulating Rns activity

The results of this study clearly delineate the role of the binding pocket in regulating the AraC-VR response to fatty acids. Charged amino acid side chains, specifically H20 and R75, stabilize the carboxylate head group of the decanoic acid and are required for full ligand binding and inhibitory activities. Interestingly, the effects of mutations varied. Removing the positive charge on R75, which had the most marked phenotype, eliminated decanoic acid inhibition in both transcriptional and DNA binding assays. However, removing the charge on H20 led to a more pronounced insensitivity to decanoic acid by EMSA than that observed in a Lac reporter strain. These differences could be the result of assay conditions, with one *in situ* and the other *in vitro*. In addition to the assays above all the structures of the electrostatic mutants were devoid of ligand. Therefore, the charged amino acids are important for FAs to bind and inhibit Rns activity.

While the overall structures of the altered Rns proteins are similar, our results indicate that variants that do not bind ligand are more flexible than the wildtype protein. In particular, the transition from an ordered helix in α2 to a disordered chain in the I14F/I17F variant is intriguingly similar to what has been observed in the ligand-free structure of *V. cholerae* ToxT (Lowden et al. 2009).

Taken together, the results from this study and previous studies support a model in which virulence gene regulators of the AraC-family utilize a common regulatory mechanism in which fatty acid binding switches the proteins from a relaxed conformation to a rigid conformation that inhibits dimerization and/or DNA binding, thereby blocking the production of virulence genes. If this model indeed proves to be true across a wide group of enteric pathogens, the potential for developing broad spectrum small molecule antivirulence therapeutics seems high.

## Materials and Methods

### Plasmid Site-Directed Mutagenesis

Plasmid pGPMRns-Myc was PCR amplified with primers 1565/1566, 1567/1568, and 2410/2411 to generate *rns* H20A, R75A, and I14F/I17F, respectively. The PCR products were circularized with NEB HiFi to construct pGPMRns-Myc H20A, R75A, H20A/R75A, and I14F/I17F for use in β-galactosidase assays.

Rns expression plasmids for protein expression were constructed by PCR amplification of pCDB24-Rns (Midgett et al. 2021) with point mutation primers designed using SnapGene followed by ligation with the KLD kit (NEB). The resulting plasmids were transformed into DH5α’s, plated onto LB agar with 100 µg/ml carbenicillin and incubated overnight at 37 °C. Selected colonies were used to inoculate ZYP-0.8G, 100 µg/ml carbenicillin and grown overnight at 37 °C. Plasmids were isolated using the Qiagen miniprep kit and the mutations were verified by sequencing. Sequence verified plasmids were then transformed into BL21 DE3 cells (NEB). Oligonucleotides used in this study are listed in (Table S1).

### MBP-Rns expression plasmids

Codon optimized Rns was amplified with primers 2469/2497 from pCDB24-Rns (WT) (Midgett et al. 2021), pJDT037 (H20A), pJDT011 (R75A), pJDT039 (H20A/R75A), or pJDT104 (I14F/I17F). The PCR products and pMalC2 were digested with BamHI and HindIII then ligated resulting in pMBPRnsOPT2 WT, H20A, R75A, H20A/R75A, and I14F/I17F, respectively. All plasmid sequences were confirmed via Sanger sequencing. Plasmids used in this study are listed in (Table S2).

### β-galactosidase assays

Lac reporter strain GPM1072 was transformed with pGPMRns-Myc plasmids (WT, H20A, R75A, H20A/R75A, I14F/I17F) or vector pTags. All strains were grown aerobically at 37 °C to stationary phase in LB medium with 100 µg/ml ampicillin with or without decanoic acid in 0.4% vol/vol DMSO. β-galactosidase activity was assayed as previously described (Miller 1972). All strains used in this study are listed in (Table S3)

### Rns variant crystallization

The Rns variants were cultured and purified as described in (Midgett et. al. 2021) with the following changes (Midgett et al. 2021)Lysis was performed with sonication on ice at 50% power for 7 minutes, cycling on for 30 seconds and off for 30 seconds. The remainder of the purification was the same.

Initial crystal trays were setup using 96 well evolution plates. The condition described in Midgett et. al. 2021 (Midgett et al. 2021) was used as a starting point. While other matrix screens from Qiagen or Hampton were tried, the original condition was found to produce the most consistent crystals of the Rns variants. Interestingly, a round of screening with the additive screen revealed that DMSO improved crystal morphology, for a final crystal condition of 0.1 M succinic acid, 14% PEG 3350, 0.03 M glycyl-glycyl-glycine. The cryogenic condition used for all crystals was 1:1 glycerol and reservoir solution resulting in 50% glycerol cryo-conditions

### Rns variant data collection, structural determination, and analysis

Diffraction data of the Rns variants was collected at NSLS2 using either the AMX or FMX beamlines. Initial data reduction was performed with XDS (Kabsch 2010; Midgett et al. 2021). Molecular replacement was performed using PHASER with the wildtype Rns PDB:6XIU as the model (Midgett et al. 2021). Automated refinement was carried out in PHENIX and manual model building was done using COOT (Adams et al. 2010; Emsley et al. 2010). Final models were deposited into the PDB (Berman et al. 2003). See supplementary table for data and model statistics (Table S4).

ChimeraX was used for structural analysis and visualization (Goddard et al. 2018). To analyze the structures, they were first aligned in Matchmaker (Meng et al. 2006) using the long helix in the DNA binding domain (N204-E223) as a fiducial marker. The opening between the N- and C-terminal domain was measured by selecting the Cα of the amino acids 73 and 219 and then using ChimeraX to obtain the distances. To obtain the overall Cα RMSD of the structures the Cα atoms were selected and then aligned in Matchmaker, each chain was aligned separately as shown in Figure S1.

To compare the B-factors of the different structures it was first necessary to scale the B-factors to a common median (Carugo 2022). To scale the B-factors of the pdb’s of the structures were edited so only the lines describing protein atom positions were left, i.e. all water, and heteroatoms were removed. The data was then imported into STATA (STATACorp) so all the pdb’s were in one dataset. Scaling was carried out using a robust scaling method, so the median B-factor was set to 0 and the interquartile range was set to 1 for each structure. The results were graphed using STATA.

### Expression and purification of MBP-Rns

Maltose binding protein (MBP) fused to the amino terminus of Rns was expressed and purified with a protocol modified from Bodero et. al., 2007 (Bodero et al. 2007). BL21 cells were transformed with pMBPRnsOpt2 WT, H20A, R75A, or I14F/I17F and grown aerobically at 37 °C in LB medium with 200 µg/ml ampicillin and 2 mM MgCl_2_. After reaching mid-log phase cultures were transferred to a 30 °C shaking water bath and expression of MBP-Rns was induced by the addition of 1 mM IPTG. After overnight induction, cells were harvested at 6,000 × *g* for 10 min at 4 °C and concentrated 100-fold in ice-cold lysis buffer A (20 mM Tris-Cl pH 7.6, 200 mM NaCl, 1 mM EDTA). Cells were lysed by two-three passages through a French press. Insoluble material was removed by centrifugation at 40,000 ×*g* for 40 min at 4 °C. The supernatant was loaded onto a 1 ml MBPTrap HP column (Cytiva) and eluted from the column with buffer A with 10 mM maltose. Fractions containing MBP-Rns were loaded onto a HiPrep Heparin FF 16/10 column (Cytiva) in heparin buffer (20 mM Tris-Cl pH 8, 150 mM NaCl) and eluted via a step gradient with heparin buffer with 1 M NaCl. Fractions containing MBP-Rns were loaded onto a Pall Centrifuge filter for buffer exchange to storage buffer (10 mM Tris-Cl pH 7.4, 280 mM NaCl, 1 mM EDTA, 10 mM β-mercaptoethanol, 25% glycerol vol/vol) and stored at -80 °C. Protein concentration was determined with the Bio-Rad Bradford assay relative to a BSA standard curve.

### Electrophoretic mobility shift assays

Oligonucleotides 2107/2108 which encode a prototypical Rns binding site that is 5’ cyanine 5.5 tagged on both strands were duplexed by heating at 65 °C for 10 mins followed by slow cooling to room temperature over greater than one hour. 30 nM purified MBP-Rns was incubated with decanoic acid (final concentration 2% DMSO vol/vol) in binding buffer (10 mM Tris-Cl pH 7.4, 50 mM KCl, 1 mM DTT, 1 ng/µl poly(dI-dC), and 100 µg/ml BSA) for one hour at room temperature. 5 nM duplex probe was added to each reaction followed by an additional incubation at room temperature for 10 mins. Glycerol was added to a final concentration of 6.5% (vol/vol) prior to separation on 5% native polyacrylamide gels. Gels were analyzed with the Odyssey FC Imaging System (LI-COR Biosciences).

#### Data sharing

The Rns variant structures were deposited into the PDB; RnsH20A (PDBID: 9CA5), RnsR75A (PDBID: 8VRQ), RnsH20A/R75A (PDBID: 8VST), and RnsI14F/I17F (PDBID: 9CA6).

## Supplemental Information

**Figure S1.**
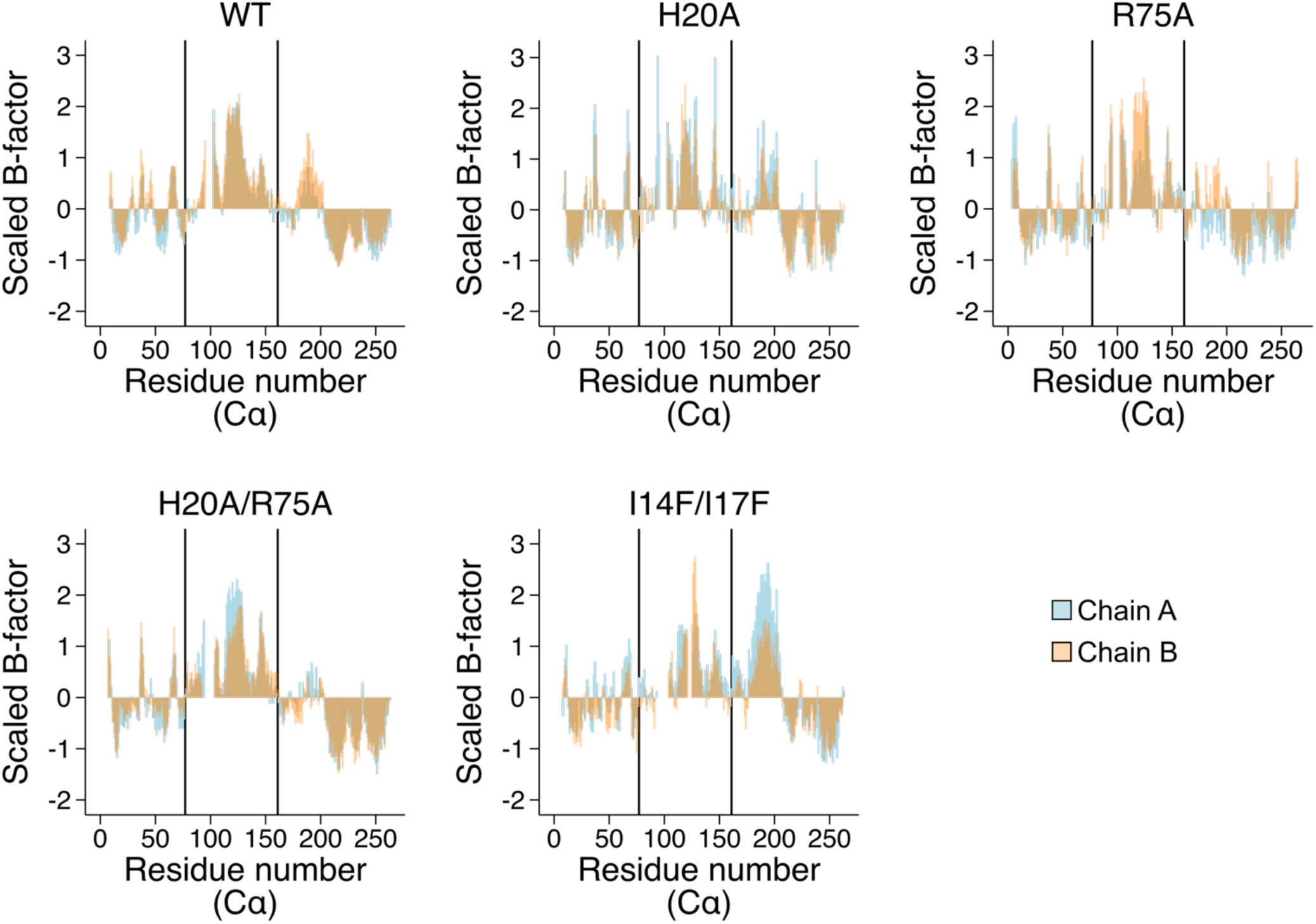
Scaled B-factors of Rns wildtype and altered protein structures. Graphs of the scaled B-factors from the different structures are shown. There are two lines drawn at residue 77 and residue 161 on each plot, separating them into three regions. From left to right the first region is most of the binding pocket, followed by the dimerization helices, and finally the DNA binding domain.

**Table S1:**
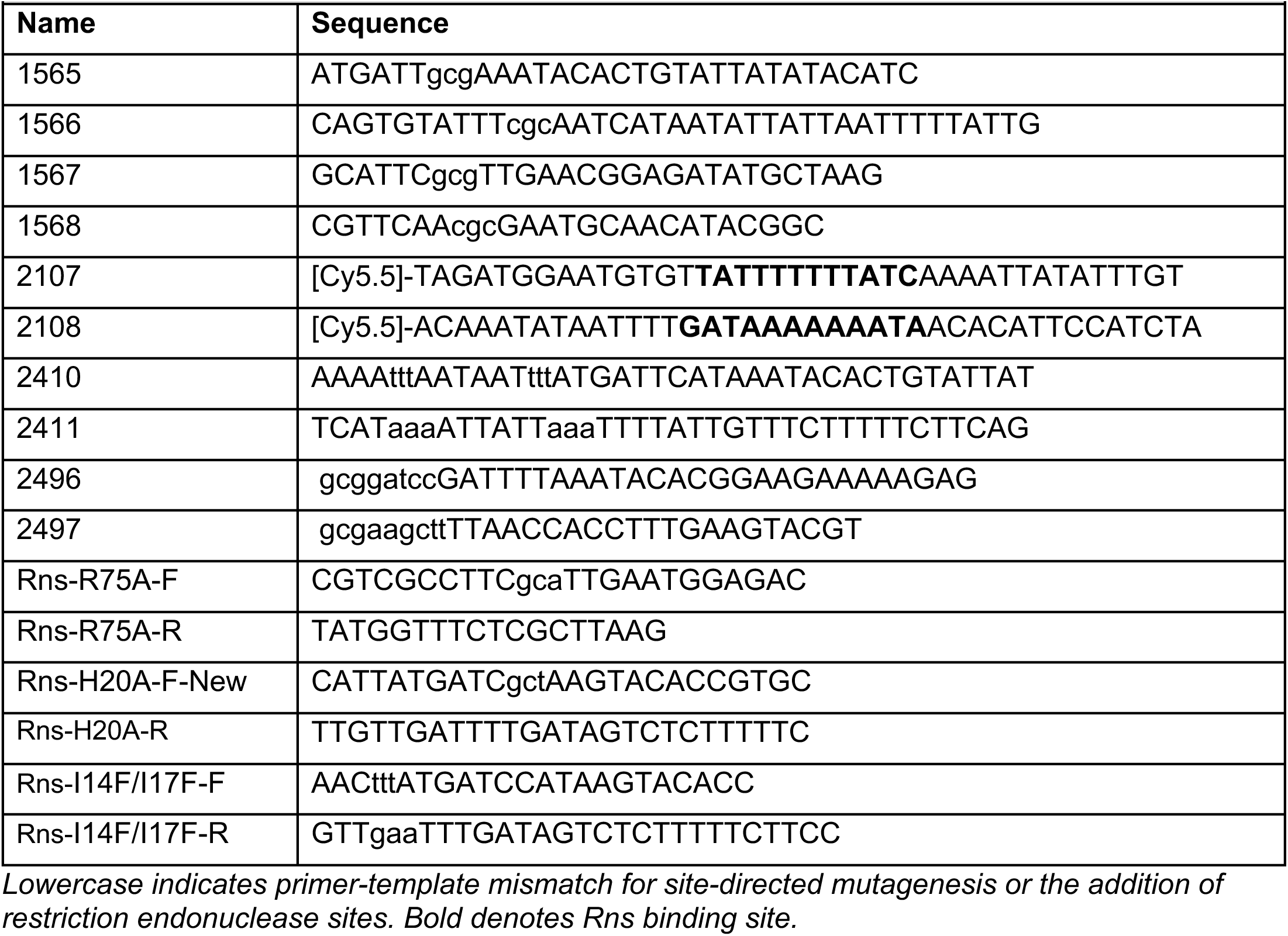
Primers used in this study.

**Table 2:** Plasmids used in this study.

| Name | Description | Marker | Reference |
| --- | --- | --- | --- |
| pGPMRns-Myc | Rns expressed from *lacp* | *bla* | [1] |
| pGPMRns-Myc H20A | Rns H20A expressed from *lacp* | *bla* | This study |
| pGPMRns-Myc R75A | Rns R75A expressed from *lacp* | *bla* | This study |
| pGPMRns-Myc H20A/R75A | Rns H20A/R75A expressed from *lacp* | *bla* | This study |
| pGPMRns-Myc I14F/I17F | Rns I14F/I17F expressed from *lacp* | *bla* | This study |
| pTags2 | Cloning vector | *bla* | [2] |
| pMal-C2 | MBP cloning vector |  | New England Biolabs |
| pCDB24-Rns | 10xHis-SUMO(SMT)-Rns |  | [1] |
| pJDT037 | SMT-Rns-H20A |  | This study |
| pJDT011 | SMT-Rns-R75A |  | This study |
| pJDT039 | SMT-Rns-H20A/R75A |  | This study |
| pJDT104 | SMT-Rns-I14F/I17F |  | This study |
| pMBPRnsOpt2 | IPTG inducible codon optimized MBP-Rns from tacp |  | This study |

**Table S3:**
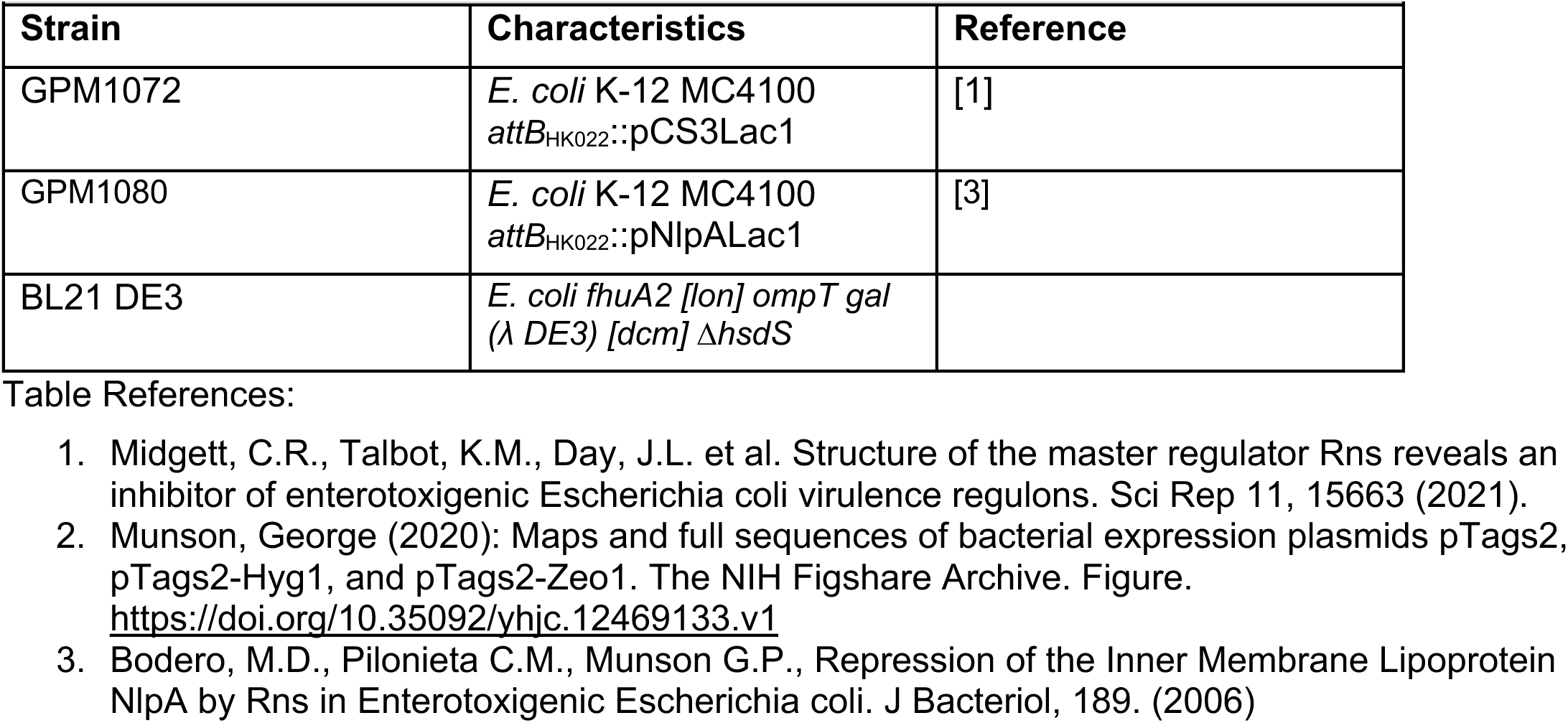
Strains used in this study.

**Table S4:** Crystallographic Statistics for all mutants.

|  | RnsR75A<br>(8VRQ) | Rns H20A/R75A<br>(8VST) | Rns I14FI17F<br>(9CA6) | H20A<br>(9CA5) |
| --- | --- | --- | --- | --- |
| Resolution range | 29.22 - 2.4<br>(2.486 - 2.4) | 48.65 - 2.4<br>(2.486 - 2.4) | 29.2 - 2.995<br>(3.1 - 3.0) | 46.1 - 2.9<br>(3 - 2.9) |
| Space group | P 21 21 21 | P 21 21 21 | P 21 21 21 | P 21 21 21 |
| Unit cell | 47.73 96.97 136.59<br>90 90 90 | 47.65 97.29 136.94<br>90 90 90 | 47.81 96.68 138.15<br>90 90 90 | 47.73 97.93 136.69<br>90 90 90 |
| Total reflections | 161945 (15846) | 170086 (17399) | 68908 (6923) | 90422 (9910) |
| Unique reflections | 25501 (2497) | 25620 (2499) | 24421 (2403) | 25426 (2759) |
| Multiplicity | 6.4 (6.3) | 6.6 (7.0) | 2.8 (2.9) | 3.6 (3.6) |
| Completeness (%) | 99.55 (99.64) | 99.70 (99.84) | 98.98 (98.91) | 92.35 (99.18) |
| Mean I/sigma(I) | 7.03 (0.74) | 6.49 (0.78) | 2.32 (0.44) | 3.08 (0.38) |
| Wilson B-factor | 63.73 | 60.41 | 77.99 | 87.44 |
| R-merge | 0.1517 (2.973) | 0.1837 (2.423) | 0.3782 (2.264) | 0.2496 (3.402) |
| R-meas | 0.1659 (3.239) | 0.1998 (2.617) | 0.4687 (2.746) | 0.295 (4.008) |
| R-pim | 0.06641 (1.273) | 0.07736 (0.9801) | 0.2723 (1.533) | 0.1558 (2.099) |
| CC1/2 | 0.996 (0.395) | 0.995 (0.383) | 0.928 (0.146) | 0.986 (0.163) |
| CC* | 0.999 (0.752) | 0.999 (0.744) | 0.981 (0.506) | 0.996 (0.529) |
| Reflections used in refinement | 25457 (2489) | 25595 (2497) | 13327 (1272) | 13705 (1448) |
| Reflections used for R-free | 1997 (196) | 2000 (195) | 1333 (128) | 1372 (146) |
| R-work | 0.2383 (0.3223) | 0.2467 (0.3711) | 0.3488 (0.4226) | 0.2641 (0.4099) |
| R-free | 0.2869 (0.3469) | 0.2971(0.3853) | 0.3678 (0.4689) | 0.3149 (0.4560) |
| Number of non-hydrogen atoms | 4151 | 4026 | 3939 | 4025 |
| macromolecules | 4137 | 4011 | 3939 | 4025 |
| Protein residues | 508 | 493 | 481 | 493 |
| RMS(bonds) | 0.01 | 0.01 | 0.009 | 0.006 |
| RMS(angles) | 1.13 | 1.18 | 1.09 | 0.75 |
| Ramachandran favored (%) | 97.6 | 98.76 | 96.79 | 96.91 |
| Ramachandran allowed (%) | 2.4 | 1.24 | 3.21 | 2.89 |
| Ramachandran outliers (%) | 0 | 0 | 0 | 0.21 |
| **Rotamer outliers (%)** | 1.27 | 2.62 | 2 | 1.3 |
| **Clashscore** | 8.72 | 10.56 | 8.91 | 8.31 |
| **Average B-factor** | 70.89 | 69.54 | 82.21 | 101.08 |

